# Novel Variants in COL4A3 and COL4A4 are Causes of Alport Syndrome in Rio Grande do Norte, Brazil

**DOI:** 10.1101/2019.12.17.878918

**Authors:** Washington Candeia de Araújo, Raul Maia Falcão, Raquel Uchoa, Carlos Alexandre Garcia, Jorge Estefano S. de Souza, Selma M. B. Jeronimo

## Abstract

**Background:** Alport syndrome is a progressive and hereditary nephropathy, characterized by hematuria and proteinuria, and extrarenal manifestations as hearing loss and eye abnormalities. The disease can be expressed as autosomal recessive or dominant, caused by variants in COL4A3 and COL4A4 *loci*, respectively, or X-linked caused by variants in COL4A5 *locus*.

**Methods:** Two unrelated families with Alport Syndrome from northeast of Brazil were studied and whole exome sequencing were performed. DNA sequences were mapped against the human genome (GRCh38/hg38 build) to identify associated mutations.

**Results:** Variant analysis showed deleterious variants in COL4A3 and COL4A4 *loci* in chromosome 2. Two variants were detected with alternative alleles in a homozygous state in the probands. One novel premature stop codon at position 481 of COL4A3 *protein* is present in one family and one frameshift mutation leading to a premature stop codon at position 786 of COL4A4 protein in the other family. Both Alport cases presented their variants surrounded by a broad runs of homozygosity (ROH).

**Conclusions:** The autosome recessive inheritance coupled with the runs of homozygosity in both families suggest inbreeding.

## Introduction

Alport syndrome is a hereditary Mendelian disease in which individuals progress with hematuric nephritis and sensorineural deafness. These symptoms are the results of defect in the glomerular basement membranes biogenesis [1]. The pathophysiology of Alport Syndrome consists of structural defects in type IV collagen alpha chains that constitutes the glomerular basement membrane [2–3]. Collagen type IV are encoded by six distinct genes (COL4A1 to COL4A6) consisting of six alpha chains (α1-α6) which assemble in heterotrimers (three α chains coiled together) forming a protomer [4–5]. There are three major mutations in α3, α4 or α5 generating defective type IV collagen networks and preventing proper structure of glomerular basement membrane [6–7].

Although there are six alpha chains, only three different protomers are known, α1α1α2 (IV), α5α5α6 (IV) and α3α4α5 (IV) [8–10]. Those protomers form the type IV collagen networks which are an essential part of the extracellular matrix and the structure of all basement membrane. Type IV protomers consists in a triple collagenous helix domain of about 400nm, characterized by a triplet amino acid repeat sequence of Gly-X-Y. Furthermore, a globular non-collagenous domain is found in the 3’ as well as a short domain, called 7S in the 5’ end. These domains form a dense collagens IV network, once secreted in the extracellular matrix [11–12].

The α1α1α2 (IV) protomer is ubiquitously distributed in all basement membranes during embryogenesis, later are replacing by the protomer α3α4α5 (IV) in glomerular basement membrane and protomer α5α5α6 in other tissues [13]. The α3α4α5 (IV) is known to be more cross-linked protomer [14–15], resulting in a stronger and more resistant heterotrimer [16] Collagen IV network has a crucial role for development of glomerular basement membrane during ontogenesis, which helps to support tissue integrity besides cell signaling, morphogenesis and tissue regeneration [4]. There is an interaction in the biogenesis of collagen produced by podocytes and the glomerular basement membrane. Moreover, α3α4α5 (IV) protomer is also produced in other tissues, such as the cochlea and eyes [17–18].

There are several variants described for the COL4A3 (l120070, OMIM) loci and COL4A4 (120131, OMIM) *loci*, localized in 2q35-37, and COL4A5 (303630, OMIM) *locus*. Autosomal recessive Alport Syndrome is related to modifications in the α3 (IV) and α4 (IV) chains, whereas X-linked Alport Syndrome is related to modifications in the α5 (IV). In the case of α6 chain, although the α5α5α6(IV) protomers can be formed in autosomal recessive Alport Syndrome. Experimental study demonstrated that this gene is not associated with Alport Syndrome [19]. Both modifications affect the biogenesis of type IV collagen and lead to a reduction or absence of these molecules in glomerular basement membrane progressing to a renal failure [20].

Inbreeding can increase the risk of Alport syndrome [21]. Recent studies have showm an increase of genomic homozygosity with increasing consanguinity [22–26]. In this work, we report the results of Alport syndrome cases from two families from the Northeast region of Brazil. We identify two novel variants (stop codon gained and frameshift) and the mutations result in a truncated α3 and α4 chains. In addition, there is a broad region of runs of homozygosity (ROH).

## Material and Methods

### Biological material from patients (Families Sampling)

Two unrelated families, Family 1 (F1) and Family 2 (F2), of the Northeast region in Brazil (Rio Grande do Norte state) were evaluated. Nuclear Family 1 had 4 cases of Alport, but within the extended family there were 5 other cases, all had been transplanted and Family 2 apparently had a sporadic case of Alport.

### Ethical Considerations

The study protocol was reviewed and approved by the Federal University Ethical Committee (CEP-UFRN 50-01) and by the Brazilian Ethical Committee (CONEP-4569). All participants and-or legal guardians read, approved and signed the informed consent. Subjects with medical conditions discovered during the study were treated by the study team or referred or taken to the appropriate medical resource in Natal. Subjects with Alport syndrome had already been transplanted at the time of recruitment and they were been followed at the University Hospital for their medical care.

### DNA extraction

DNA was extracted from 10 ml of anticoagulated whole blood (EDTA) by erythrocyte lysis in 70 μg of NH_4_H_2_CO_3_/ml and 7.0 mg of NH_4_Cl/ml followed by lysis of leukocytes in 1% sodium dodecyl sulfate, 100 mM EDTA, and 200 mM Tris (pH 8.5) and precipitation in isopropanol. DNA sample underwent quality control which included gel electrophoresis and quantification and determination of integrity by fluorometer (qubit, Themo Fisher, USA).

### Whole Exome Sequencing

In each family the father, mother, unaffected sibling and proband (n= 4) had their exomes sequenced. Whole exome sequencing (WES) was performed by the University of Iowa Genomics Division using manufacturer recommended protocols. Briefly, 3µg of genomic DNA were sheared using the Covaris E220 sonicator. The sheared DNA was used to prepare indexed whole exome sequencing libraries using the Agilent SureSelect XT Human all exon v6 + UTR kit (Agilent Technologies, Santa Clara, CA, Cat. No. 5190-8883). The molar concentrations of the indexed libraries were measured using the Fragment Analyzer (Agilent Technologies) and combined equally into pools for sequencing. The pooled libraries were loaded on an Illumina HiSeq 4000 genome sequencer using the 2 x 150bp paired-end sequencing by synthesis (SBS) chemistry.

### Raw Reads Preprocessing and Reference Genome Mapping

FastQC (v0.11.4) [27] software was used to quantify the sequences and to analyze the sequencing quality. Adapters was removed by Trim Galore (v0.4.1) [28] using the option to execute Cutadapt (v1.8.3) [29], with default values. Reads were mapped using BWA (v0.7.12-r1039) [30] software, with mem mode against to GRCh38/hg38 build [31]. Functions stats and depth from SAMtools (v1.7) [32] were used to calculate total reads mapped, total exome mean coverage and mean coverage in ROH and remaining regions. Normalized mean coverage was calculated dividing the average coverage per base by the individual number of reads sequenced then multiplying by the average of reads sequenced of all individuals. Also, mpileup function from SAMtools was used to generate input to VarScan2 (v2.3.9) [33] software.

### Runs of Homozygosity (ROH) ratio

To obtain total chromosomal ROH lengths the algorithm H3M2 [34] was used followed by a filter of long ROH fragments lengths (1.5Mb). The software was applied in the mapping file for each individual sample of this study. A reference length of ROH was determined by applying H3M2 with the same filter in 20 random publicly available CEU samples sequenced by the 1000 Genomes Project [35]. Thus, as a measure of the consanguinity/endogamy a homozygous ratio was calculated dividing the total ROH length of each family member by the average of ROH length from CEU samples.

### Variant Calling and Data Refinement

Best practices from Genome Analysis ToolKit (v3.4-46) HaplotypeCaller [36] (GATK-HC) were performed with *Base Quality Score Recalibration* (BQSR) mode using the following known sites: The Single Nucleotide Polymorphism Database (dbSNP 146); Mills & 1000G gold standard indels for hg38 and 1000G phase1 SNPS high confidence sites for hg38. Vcftools (v0.1.13) [37] was used to filter low coverage variants and to maintain records with at least one sample with variant call data. Variants with minDP < 20 and max-missing > 0.125 were eliminated from the analysis. Vcfstats [38] from vcflib package was used to summarize basic variant calling statistic.

### Homozygosity Mapping, Variant Annotation and Haplotype Phasing

To detect differences between the homozygous and heterozygous signals Homozygous Stretch Identifier - HomSI (v2.1) [39] - software was used with default parameters values. Databases such as Exome Aggregation Consortium [40], 1000 Genomes Project and dbSNP 146 were integrated and variants were annotated by ANNOVAR (v2018Apr16) [41] and SnpEff (v 4.1) [42] software, respectively. Missing data inference and haplotype phasing were performed using Beagle (v5.1) [43] software.

### Identification of deleterious variants

The analysis focused on identifying homozygous pathogenic variants. We restricted the exome sequence data to variants with 1) allelic frequencies < 1% in Exome Aggregation Consortium database; 2) gene and protein expression (> 1 FPKM) in kidney, cochlea and eye from Illumina Body Map, The Human Protein Atlas (HPA) and Genotype-Tissue Expression Project database; 3) clinical significance from ClinVar database. Variants with a structural effect (frameshift and stop codon gained) were considered deleterious.

## Results

### Genotyping and exome Screening

On average 98.6% of the exome-wide was covered with at least 20-fold coverage at ~190x of coverage (Supplementary Table S1). A total of 106,726 (97,327 SNVs, 9 MNPs and 9,390 INDELS) variants were reported and Ti/Tv ratio was equal to 2.4. From those, 16,830 variants were nonsynonymous, 124 stop codons gained and 262 were frameshifts.

A total of 12 deleterious variants were identified (1 stop codon gained and 11 frameshift) with alternative allele homozygous state exclusively in probands (Supplementary Table S2). From these 12 variants, 8 variants had an allele frequency greater than 1% in the Exome Aggregation Consortium database including one variant classified as benign clinical significance in the clinvar database; 2 variants were not expressed in any of the organs involved in Alport disease. Two of the 12 variants were classified as possibly pathogenic and present in genes expressed in the kidney, eye and cochlea. The two later variants are directly linked to collagen type IV production and present in COL4A3 (p.Try481*) and COL4A4 (p.Val741fs) *loci*. No deleterious variants were found in COL4A5.

### Variants along COL4A genes

Twenty-three variants (Supplementary Table S3) were present in COL4A genes. Twelve variants were presents in COL4A4 (6 synonymous, 5 non-synonymous and 1 frameshift) and eleven in COL4A3 (3 synonymous, 7 non-synonymous and 1 stop gain). Figure 1 shows the distribution of all nonsynonymous and deleterious variants in this study in COL4A4, COL4A3 and COL4A5 *loci*. Only stop gained and frameshift variants were not reported in the ClinVar database. All other variants were reported on Clinvar as benign for cases of Alport Syndrome.

**Figure 1.**
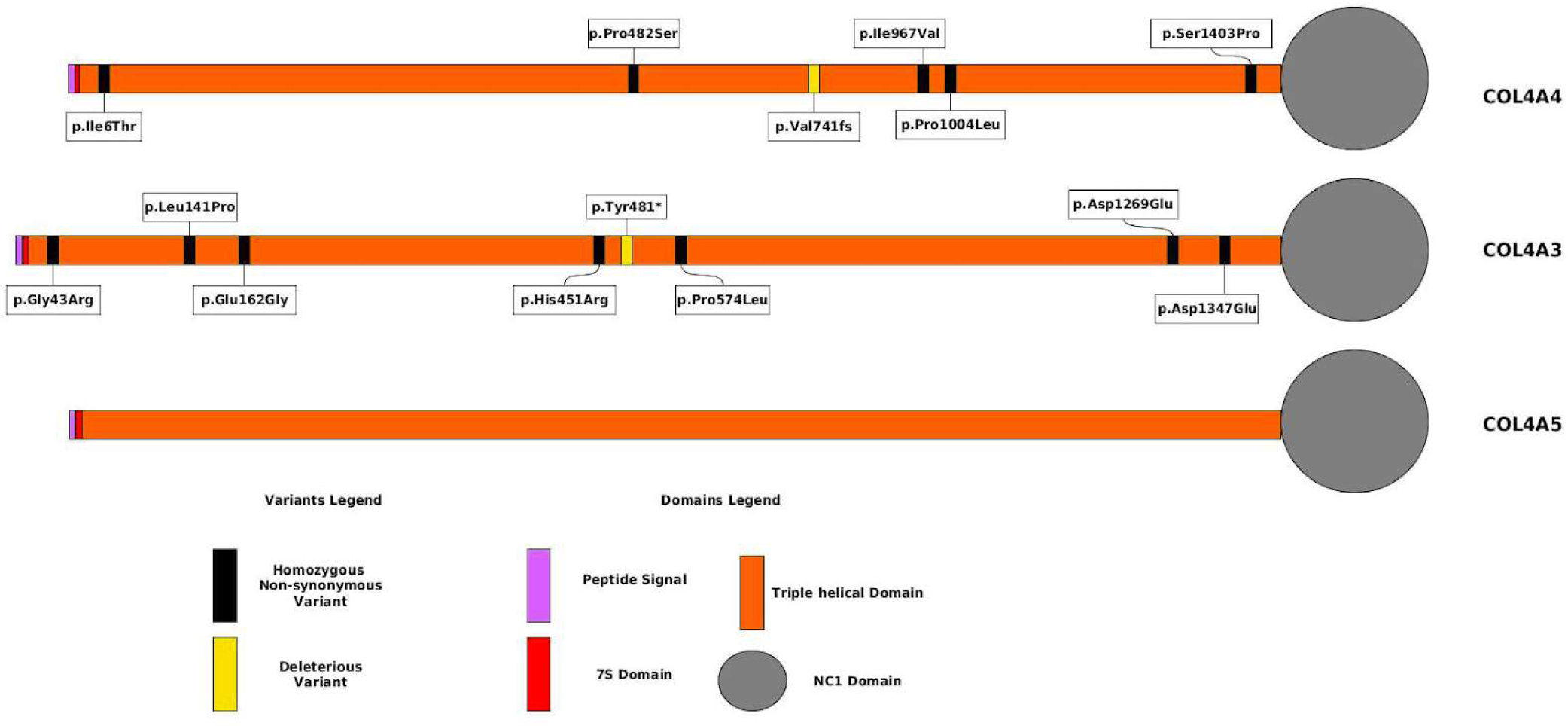
Distribution of deleterious and homozygous non-synonymous variants in COL4A4, COL4A3 and COL4A5 genes.

Eleven variants in COL4A3 occurred at triple helix region domain of collagen type IV. We observed a premature stop codon gained (p.Try481*) in all members among Family 1. In this family, proband presented this variant with alternative allele in homozygous state. Other individuals of this family had the same variant at heterozygous state. All members of Family 1, except the proband, have the same genotype in this locus, whereas members of Family 2 do not have this variant.

Nine variants in COL4A4 occurred at triple helix region domain of collagen type IV. We observed a frameshift (p.Val741fs) in all members among Family 2. In this family, proband had a homozygous frameshift leading to a truncated protein with 786 amino acids. Other members of this family had the same frameshift at heterozygous state. Members of Family 1 do not have this variant.

### Segregation analysis in COL4A genes

Haplotype phasing showed offspring’s alleles were inherited together from copies of paternal and maternal chromosomes. Both probands presented an excess of nonsynonymous variants at homozygous state. For Family 1, proband had four variants in COL4A3 and all remain members had the same three variants in COL4A3. Conversely, Family 2 proband had four variants in COL4A3 and three in COL4A4 loci while all remain members had at most three variants distributed in both COL4A genes. All variants of the genes encoding α4 and α3 chains of collagen type IV perfectly co-segregated with the disease in both families (Figure 2). All individuals showed approximately 190x (Table 1) of coverage in homozygous regions and remaining regions.

**Table 1:** Normalized mean coverage by samples in ROH and remaining regions for complete chromosome 2 extension

| Sample | Total Mapped reads to chr2 | Normalized mean coverage in homozygous region | Normalized mean coverage in remaining regions |
| --- | --- | --- | --- |
| Father (F1) | 11,361,323 | 191.45 | 195.62 |
| Mother (F1) | 12,778,526 | 186.90 | 190.10 |
| Son (F1) | 14,695,917 | 183.38 | 184.14 |
| Proband (F1) | 10,429,522 | 189.51 | 191.46 |
| Father (F2) | 15,160,228 | 192.15 | 191.13 |
| Mother (F2) | 13,382,947 | 198.62 | 192.09 |
| Son (F2) | 13,375,114 | 198.19 | 194.00 |
| Proband (F2) | 11,124,247 | 200.07 | 194.61 |

**Figure 2.**
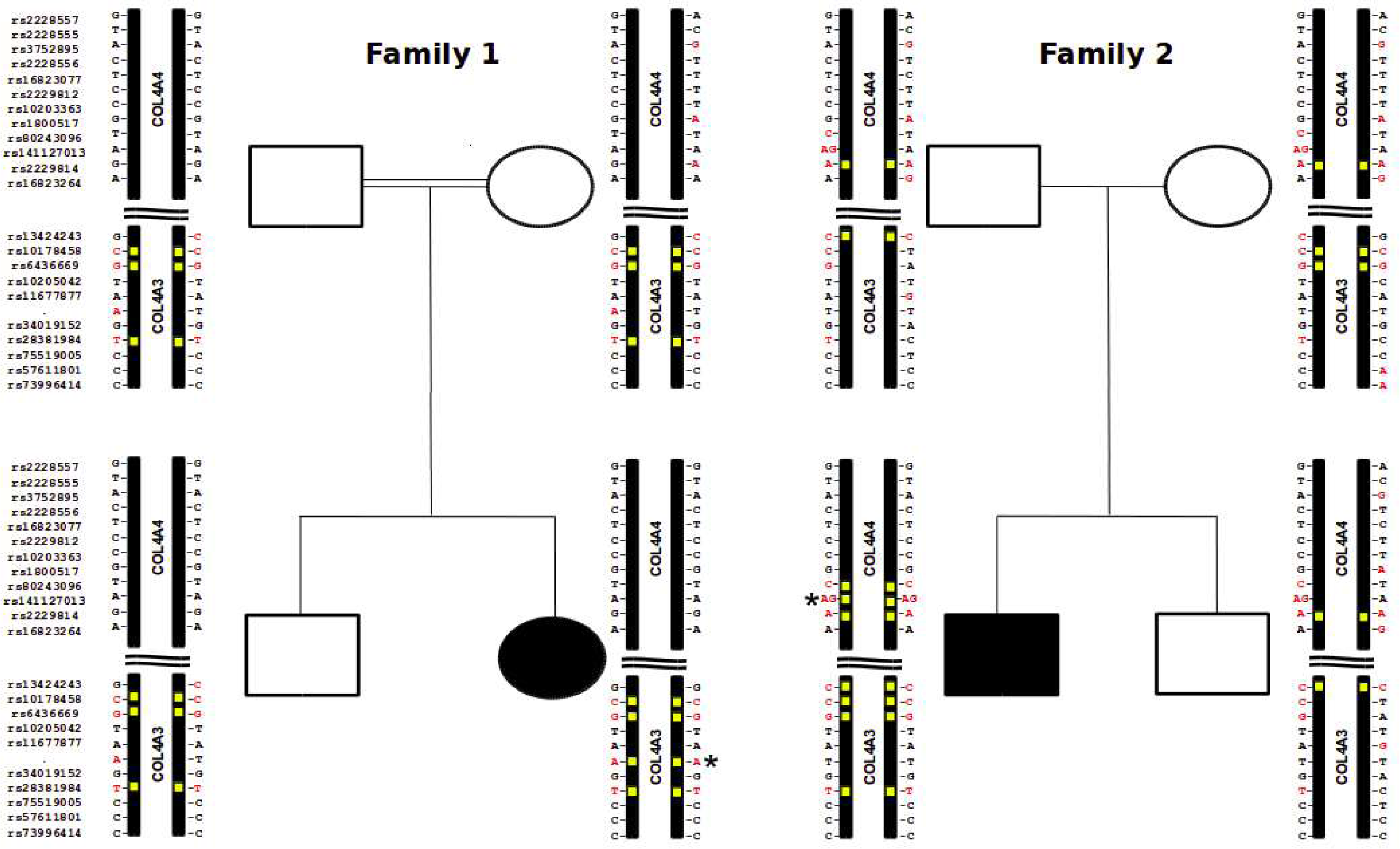
COL4A4 and COL4A3 alleles co-segregating. Figure indicates the variants of chromosome 2 for COL4A4 and COL4A3 of all individuals in families, in homozygosity (non-synonymous variant in red, homozygous non-synonymous variant in yellow). The probands are in black. Black * defines a stop codon gain in proband of Family 1 and a frame-shift in proband of Family 2.

Non-synonymous variant alleles found in COL4A genes, when analyzed individually, occurred at high allelic frequency rates in the 1KGP database. However, their co-occurrence with haplotype frequency was rare in all cases. Co-occurrence including deleterious variants presented in this study did not occur in any 1KGP individual.

### Homozygous regions in COL4A genes

Chromosome 2 showed a common ROH locus in both probands (Figure 3). In Family 1 proband has a homozygous locus of 20Mb with 140 genes. In Family 2 proband has a homozygous locus of 75Mb with 364 genes. By analyzing this region, we observed that only the COL4A3 and COL4A4 genes had deleterious variants.

**Figure 3.**
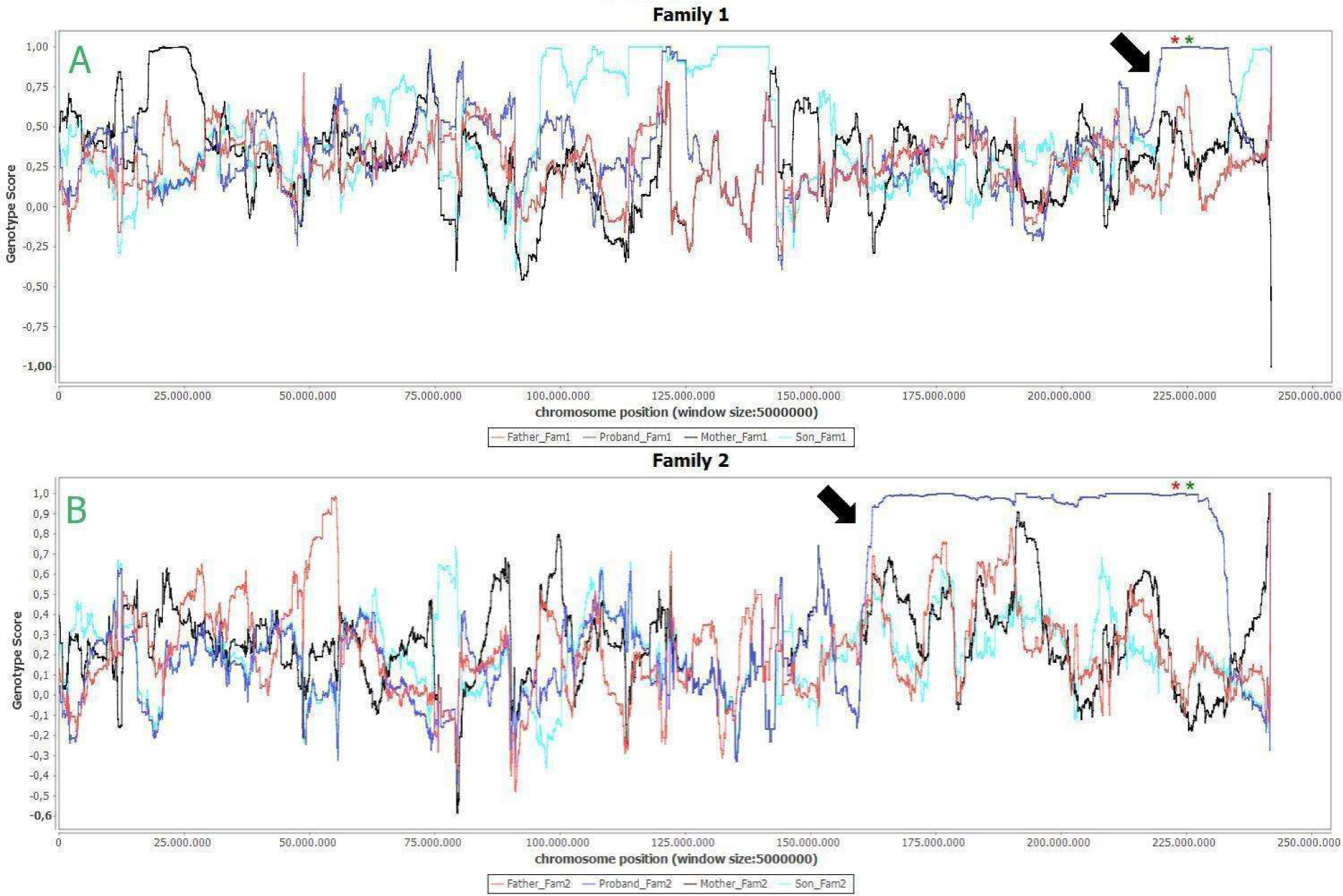
Differences between homozygous and heterozygous signals in complete chromosome 2 extension. A) Family 1, B) Family 2. Black arrow shows where homozygosity region begins in both probands (dark blue line). Red and green * define COL4A4 and COL4A3 genes respectively.

An average of 116,876,596 bp of ROH length for CEU population (Supplementary Table S4). The total of ROH fragments, ROH length for all autosomal chromosomes and homozygous ratio for each member of the family (Table 2 and 3) showed high level of consanguinity, with high homozygous ratio in the offspring. Family 1 had a homozygous ratio of 1.05, 2.15, 3.53 and 2.03 for the father, the mother, the unaffected son and the proband respectively. Family 2 had a homozygous ratio of 0.70, 0.56, 3.15 and 2.09 for father, mother, unaffected son and proband respectively. This result highlighted a relation with the number of long fragments of ROH and homozygous ratio: the higher values of fragments, the higher values of homozygous ratio.

**Table 2.** Number and total length of ROHs of the Family 1.

| **Sample** | **Number of ROHs**<br>**(cut-off 1.5 Mb)** | **Total ROH length**<br>**(cut-off 1.5 Mb)** | **Homozygous ratio** |
| --- | --- | --- | --- |
| **Father** | 44 | 123,049,309 | 1.05 |
| **Mother** | 55 | 251,514,461 | 2.15 |
| **Son** | 72 | 413,645,034 | 3.53 |
| **Proband** | 47 | 237,564,469 | 2.03 |

**Table 3.**
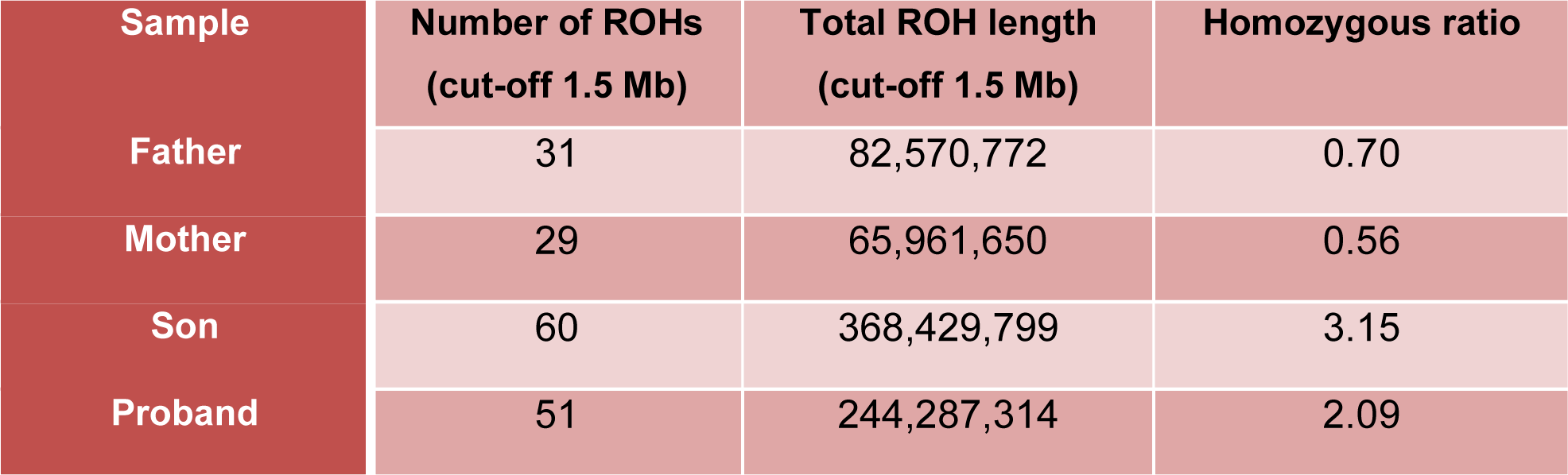
Number and total length of ROHs of the Family 2.

## Discussion

In the present study, we investigated two families from northeast of Brazil with Alport Syndrome and we report two novel variants responsible for the disease. The first one was a novel nonsense mutation in COL4A3 gene in proband from Family 1. The second one was a frameshift in COL4A4 locus in proband from Family 2. Both findings occurred in triple helix domain and had a broad region of ROH covering both genes in probands. Cosgrove and Liu [8] states that stop codon gained, frameshifts and other mutations that result in chain termination are associated with a more severe phenotype.

Szpiech et al [44] using data obtained by whole exome sequencing demonstrated that a significantly larger fraction of all predicted harmful homozygous variants are surrounded by ROH stretch when compared to those corresponding fractions of non-damaging homozygous variants. Long ROH (> 1.5 Mb) are known to be favored by inbreeding and IBD inheritance processes and characterized by an enrichment of deleterious recessive mutations [45, 46]. Most recessive disease variants are identified via homozygosity mapping. Using high-density genome-scan data in European populations McQuillan et al [24] indicate that the numbers and lengths for ROHs above 1.5Mb are clearly relation to endogamy and isolation behind the effects on ROHs, as occurred in our Alport syndrome.

A homozygous region was present in both probands in chromosome two but being broader in Family 2 when compared to Family 1 (Figure 2). Although unaffected offspring had the highest value for homozygous ratio (Table 2 and Table 3), in both families, type IV collagen is not affected. A possible explanation for this excessive homozygosity in the *loci* region of COL4A3 and COL4A4, would be a large deletion occurring in the probands leading to uniparental inheritance. However, the phasing of chromosomes as well as the average of coverage in ROH and remaining regions (Figure 2 and Table 1) pointed out to an allele inheritance and not to a deletion inheritance rejecting a hypothesis of deletion event. Recent studies [47, 48] reported that mutations in COL4A3 and COL4A4 *loci*, when in *cis*, can lead to a form of benign familial hematuria, in a similar genetic way as Alport syndrome. Benign familial hematuria is considered as a phenotypic expression and represents a carrier state for autosomal recessive Alport syndrome.

All family 1 members have a nonsense variant (p.Tyr481*) in COL4A3 gene, although only the proband has it with alternative allele at homozygous state. This variant is novel. In addition, it is known that COL4A3 nonsense phenotype is severe and associated with end-stage renal disease, deafness and ocular lesions [11]. As reported in this study, the observed stop codon gained falls in collagenous domain of type IV collagen leading to a truncated protein, and consequently destabilizing collagen IV network formation. Other mutations may increase the risk of AS because they had perfectly co-segregated with disease. In this family we observed all members having high homozygosity rate.

Members of family 2 had a frameshift variant (p.Val741fs) in COL4A4 gene, but only the proband has with alternative allele at homozygous recessive state (see Table 1). Frameshift is the most important evidence for Alport Syndrome heritage in this family due to the insertion of one base in the reading frame and led to a truncated protein with 786 residues. Thus, herein we propose Alport Syndrome resulting in phenotype of proband in this family. Zhu *et al* [49] obtained a similar result using next generation sequencing and identified a frameshift leading to truncated COL4A4 protein with 1634 amino acids and considering as a loss of function variant. In this family we noticed only the offspring had high homozygosity rate. This reflects origin from a small isolated population and presents features such as population genetic historical and geographical factors.

### Conclusions

This work describes two novel variants responsible for Alport cases in two families from northeast of Brazil. Those variants were searched in the databases of 1000 genomes and Clinvar and they are rare alleles, not yet identified in the world population. The two pathogenic variants identified are located on an ROH locus. Interestingly both Alport probands have ROH in the COL4A3 and COL4A4 loci, and all variants are homozygous. There is a need to screen people who have lost kidney function for genetic mutations, allowing a better understanding of Alport Syndrome and their respective penetrance. In addition, it is important to identify people heterozygous for variants associated with Alport Syndrome in areas with high level of inbreeding.

## Acknowledgement

The authors thank Jose Wellington for his help in recruitment of the Alport families and the staff from the Hospital Universitario Onofre Lopes. The authors also thank the Alport families for participating in this study. WCA received a fellowship from CAPES/FAPERN and RMD to CAPES and Rede de Pesquisa em Genômica Populacional Humana (RedePGH) supported by Universidade Federal do Pará (UFPA). All analysis was performed at Supercomputer of UFRN (23001011020P6 Project).

---

**Supplementary Table S1:** Exome-wide base coverage summary

| Sample | Total Mapped reads | Total exome mean coverage (normalized) | Total Covered Bases (20x fold) | % 20x fold bases covered |
| --- | --- | --- | --- | --- |
| Father (F1) | 157,896,820 | 194.15 | 89,152,408 | 98.54 |
| Mother (F1) | 180,487,311 | 189.62 | 89,312,826 | 98.71 |
| Son (F1) | 204,975,749 | 184.55 | 89,624,633 | 99.06 |
| Proband (F1) | 147,647,415 | 191.20 | 88,816,063 | 98.16 |
| Father (F2) | 212,010,435 | 191.23 | 89,699,378 | 99.14 |
| Mother (F2) | 187,140,367 | 193.47 | 89,334,742 | 98.74 |
| Son (F2) | 185,054,786 | 194.51 | 89,003,689 | 98.37 |
| Proband (F2) | 153,688,254 | 195.54 | 88,989,473 | 98.36 |

**Supplementary Table S2:**
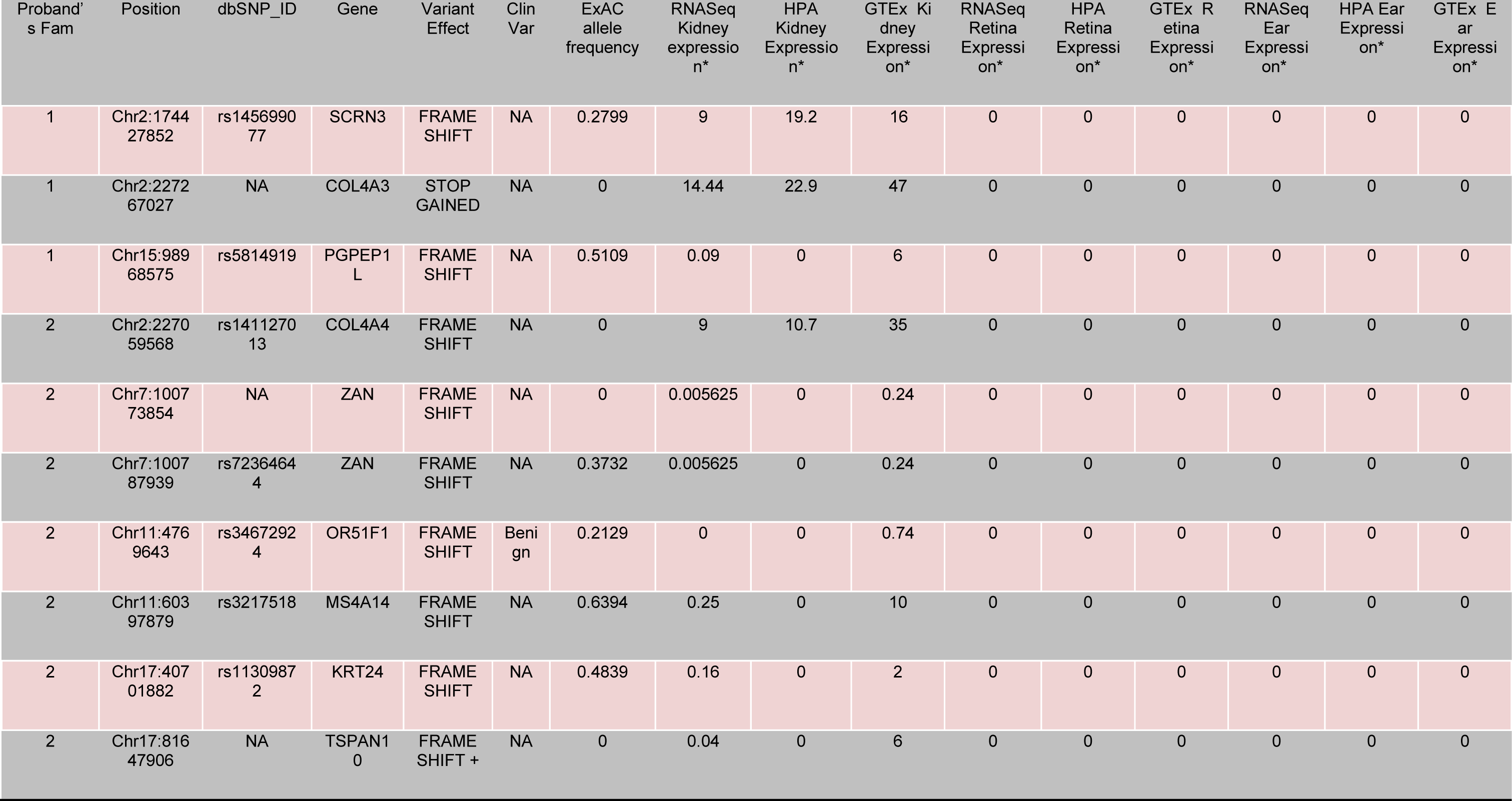
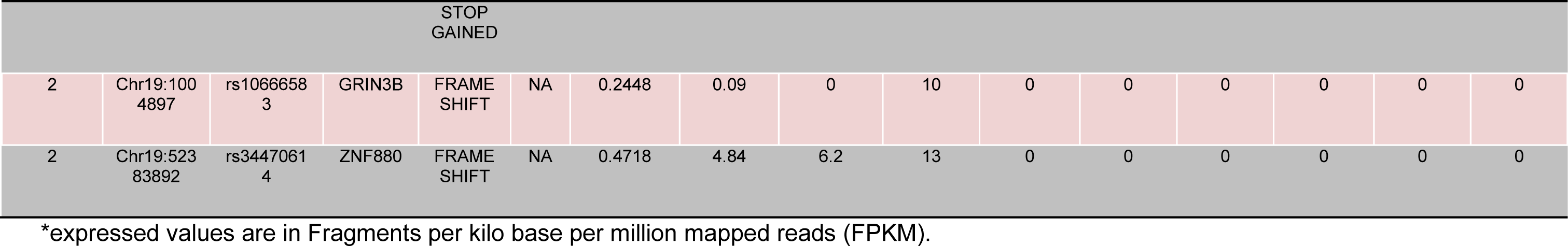
Possibly deleterious variants with alternative allele at homozygous state in probands and at heterozygous state in remains members of the family

**Supplementary Table S3:**
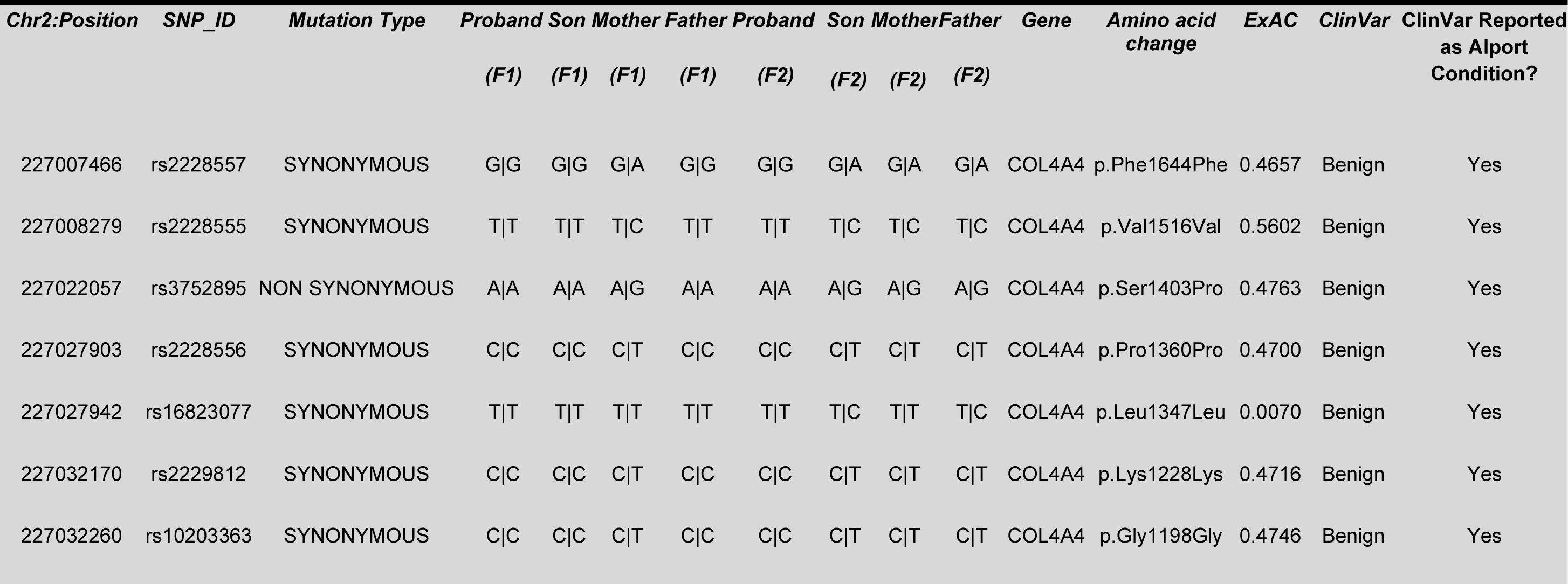
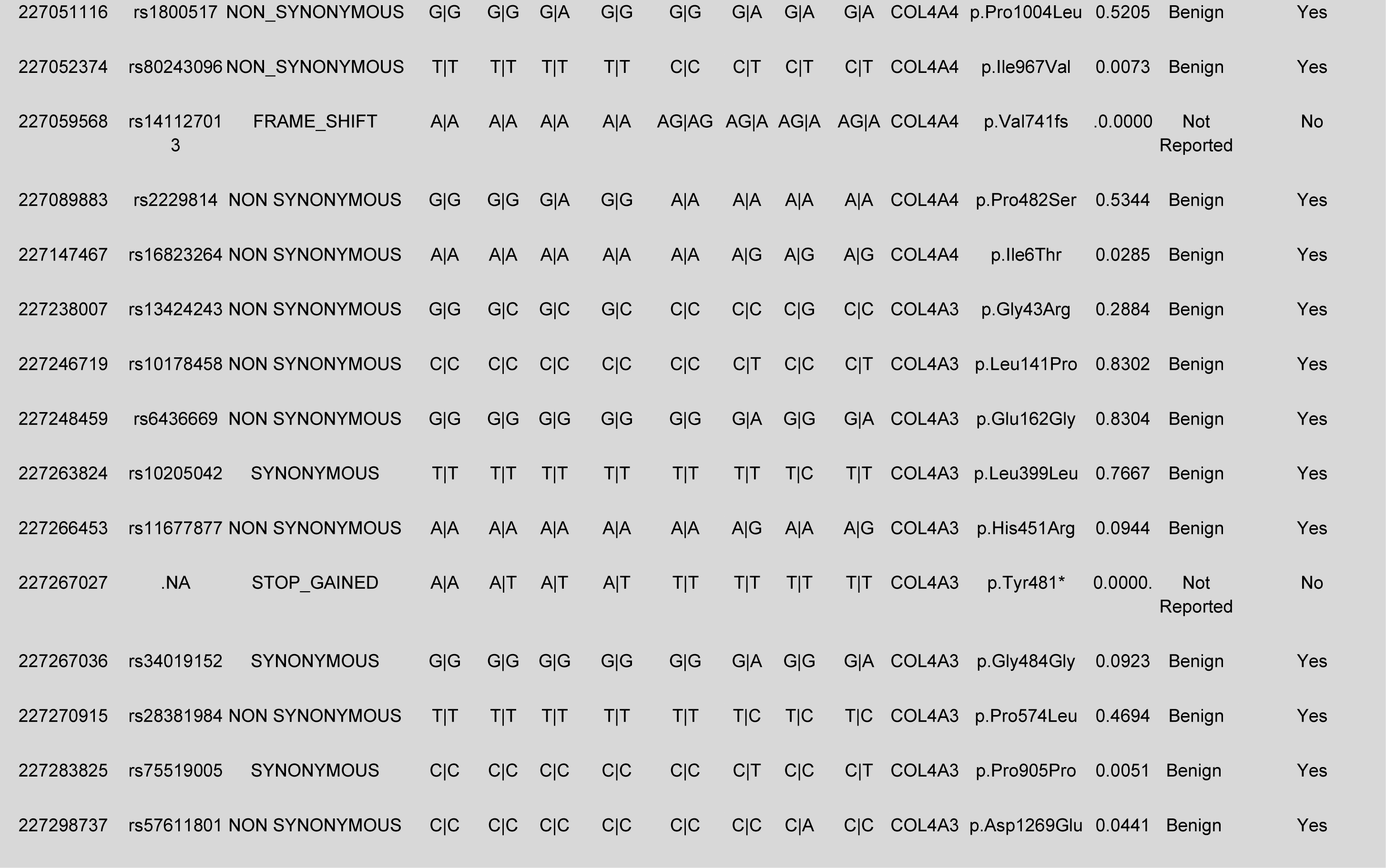
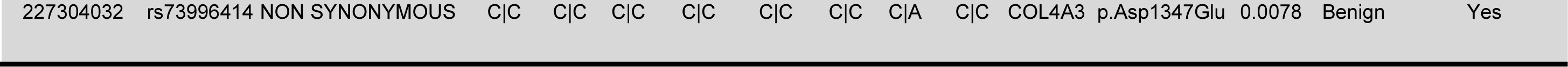
Coding variants found by NGS in COL4A4 and COL4A3 genes in chromosome 2.

| Chr2:Position | SNP_ID | Mutation Type | Proband | Son | Mother | Father | Proband | Son | Mother | Father | Gene | Amino acid change | ExAC | ClinVar | ClinVar Reported as Alport Condition? |
| --- | --- | --- | --- | --- | --- | --- | --- | --- | --- | --- | --- | --- | --- | --- | --- |
|  |  |  | (F1) | (F1) | (F1) | (F1) | (F2) | (F2) | (F2) | (F2) |  |  |  |  |  |
| 227007466 | rs2228557 | SYNONYMOUS | G\|G | G\|G | G\|A | G\|G | G\|G | G\|A | G\|A | G\|A | COL4A4 | p.Phe1644Phe | 0.4657 | Benign | Yes |
| 227008279 | rs2228555 | SYNONYMOUS | T\|T | T\|T | T\|C | T\|T | T\|T | T\|C | T\|C | T\|C | COL4A4 | p.Val1516Val | 0.5602 | Benign | Yes |
| 227022057 | rs3752895 | NON SYNONYMOUS | A\|A | A\|A | A\|G | A\|A | A\|A | A\|G | A\|G | A\|G | COL4A4 | p.Ser1403Pro | 0.4763 | Benign | Yes |
| 227027903 | rs2228556 | SYNONYMOUS | C\|C | C\|C | C\|T | C\|C | C\|C | C\|T | C\|T | C\|T | COL4A4 | p.Pro1360Pro | 0.4700 | Benign | Yes |
| 227027942 | rs16823077 | SYNONYMOUS | T\|T | T\|T | T\|T | T\|T | T\|T | T\|C | T\|T | T\|C | COL4A4 | p.Leu1347Leu | 0.0070 | Benign | Yes |
| 227032170 | rs2229812 | SYNONYMOUS | C\|C | C\|C | C\|T | C\|C | C\|C | C\|T | C\|T | C\|T | COL4A4 | p.Lys1228Lys | 0.4716 | Benign | Yes |
| 227032260 | rs10203363 | SYNONYMOUS | C\|C | C\|C | C\|T | C\|C | C\|C | C\|T | C\|T | C\|T | COL4A4 | p.Gly1198Gly | 0.4746 | Benign | Yes |
| 227051116 | rs1800517 | NON_SYNONYMOUS | G\|G | G\|G | G\|A | G\|G | G\|G | G\|A | G\|A | G\|A | COL4A4 | p.Pro1004Leu | 0.5205 | Benign | Yes |
| 227052374 | rs80243096 | NON_SYNONYMOUS | T\|T | T\|T | T\|T | T\|T | C\|C | C\|T | C\|T | C\|T | COL4A4 | p.Ile967Val | 0.0073 | Benign | Yes |
| 227059568 | rs14112701<br>3 | FRAME_SHIFT | A\|A | A\|A | A\|A | A\|A | AG\|AG | AG\|A | AG\|A | AG\|A | COL4A4 | p.Val741fs | .0.0000 | Not<br>Reported | No |
| 227089883 | rs2229814 | NON SYNONYMOUS | G\|G | G\|G | G\|A | G\|G | A\|A | A\|A | A\|A | A\|A | COL4A4 | p.Pro482Ser | 0.5344 | Benign | Yes |
| 227147467 | rs16823264 | NON SYNONYMOUS | A\|A | A\|A | A\|A | A\|A | A\|A | A\|G | A\|G | A\|G | COL4A4 | p.Ile6Thr | 0.0285 | Benign | Yes |
| 227238007 | rs13424243 | NON SYNONYMOUS | G\|G | G\|C | G\|C | G\|C | C\|C | C\|C | C\|G | C\|C | COL4A3 | p.Gly43Arg | 0.2884 | Benign | Yes |
| 227246719 | rs10178458 | NON SYNONYMOUS | C\|C | C\|C | C\|C | C\|C | C\|C | C\|T | C\|C | C\|T | COL4A3 | p.Leu141Pro | 0.8302 | Benign | Yes |
| 227248459 | rs6436669 | NON SYNONYMOUS | G\|G | G\|G | G\|G | G\|G | G\|G | G\|A | G\|G | G\|A | COL4A3 | p.Glu162Gly | 0.8304 | Benign | Yes |
| 227263824 | rs10205042 | SYNONYMOUS | T\|T | T\|T | T\|T | T\|T | T\|T | T\|T | T\|C | T\|T | COL4A3 | p.Leu399Leu | 0.7667 | Benign | Yes |
| 227266453 | rs11677877 | NON SYNONYMOUS | A\|A | A\|A | A\|A | A\|A | A\|A | A\|G | A\|A | A\|G | COL4A3 | p.His451Arg | 0.0944 | Benign | Yes |
| 227267027 | .NA | STOP_GAINED | A\|A | A\|T | A\|T | A\|T | T\|T | T\|T | T\|T | T\|T | COL4A3 | p.Tyr481* | 0.0000. | Not<br>Reported | No |
| 227267036 | rs34019152 | SYNONYMOUS | G\|G | G\|G | G\|G | G\|G | G\|G | G\|A | G\|G | G\|A | COL4A3 | p.Gly484Gly | 0.0923 | Benign | Yes |
| 227270915 | rs28381984 | NON SYNONYMOUS | T\|T | T\|T | T\|T | T\|T | T\|T | T\|C | T\|C | T\|C | COL4A3 | p.Pro574Leu | 0.4694 | Benign | Yes |
| 227283825 | rs75519005 | SYNONYMOUS | C\|C | C\|C | C\|C | C\|C | C\|C | C\|T | C\|C | C\|T | COL4A3 | p.Pro905Pro | 0.0051 | Benign | Yes |
| 227298737 | rs57611801 | NON SYNONYMOUS | C\|C | C\|C | C\|C | C\|C | C\|C | C\|C | C\|A | C\|C | COL4A3 | p.Asp1269Glu | 0.0441 | Benign | Yes |
| 227304032 | rs73996414 | NON SYNONYMOUS | C\|C | C\|C | C\|C | C\|C | C\|C | C\|C | C\|A | C\|C | COL4A3 | p.Asp1347Glu | 0.0078 | Benign | Yes |

**Supplementary Table S4:** CEU populations of the 1000 Genomes Project.

| Supplementary Table S4: CEU populations of the 1000 Genomes Project. |  |  |
| --- | --- | --- |
| CEU sample | #ROH regions (cut-off 1.5 Mb) | Total ROH length (cut-off 1.5 Mb) |
| NA07347 | 39 | 106,376,199 |
| NA11831 | 48 | 136,881,072 |
| NA11830 | 38 | 101,034,986 |
| NA10851 | 47 | 129,703,972 |
| NA11832 | 44 | 107,741,456 |
| NA07357 | 55 | 152,663,835 |
| NA11843 | 55 | 149,934,599 |
| NA06989 | 48 | 113,842,564 |
| NA07056 | 53 | 165,109,126 |
| NA06984 | 40 | 122,85,835 |
| NA07037 | 33 | 81,158,923 |
| NA06985 | 48 | 122,109,441 |
| NA07051 | 40 | 91,727,641 |
| NA07048 | 33 | 93,706,363 |
| NA11840 | 40 | 98,139,215 |
| NA11829 | 42 | 129,474,178 |
| NA10847 | 36 | 99,052,682 |
| NA07000 | 48 | 134,365,628 |
| NA06986 | 40 | 114,937,185 |
| NA06994 | 33 | 86,687,035 |

